# Fast and standardized motor-hotspot determination with automated TMS mapping

**DOI:** 10.1101/2025.09.24.678288

**Authors:** Ida Granö, Olli-Pekka Kahilakoski, Mikael Laine, Miriam Kirchhoff, Oskari Ahola, Ana M. Soto, Renan H. Matsuda, Aino E. Nieminen, Giulia Pieramico, Heikki Sinisalo, Ilkka Rissanen, Timo Tommila, Timo Roine, Victor H. Souza, Risto J. Ilmoniemi, Pantelis Lioumis, Tuomas P. Mutanen

## Abstract

Determining the optimal stimulation target for motor responses (motor hotspot) and the required intensity for reliably eliciting said responses (motor threshold) are common procedures in transcranial magnetic stimulation (TMS) research and treatments. However, the procedures for determining them are user-dependent, slow, and lack standardization, leading to long stimulation sessions with potentially inadequate outcomes. Partially automated algorithms for determining the motor threshold have been developed, but the motor hotspot is still largely mapped by hand. Automating the hotspot mapping will accelerate the process and improve standardization and accuracy. We developed a fully automated algorithm for finding the motor hotspot with multi-locus TMS and Bayesian optimization. Tested online in five healthy participants, the algorithm located motor hotspots with (mean ± 95% CI) 2.1 ± 0.7 mm and 6 ± 2° difference from the global best target with only (mean) 47 stimuli. This is a significant improvement from previous motor-mapping algorithms, which do not optimize for stimulation location and orientation simultaneously. This accurate, fast, and user-independent procedure paves the way for faster experimental processes and more streamlined clinical applications.

## Introduction

Transcranial magnetic stimulation (TMS) is a powerful tool for non-invasive cortical mapping of motor and speech functional areas (Krieg et al., 2017, 2016; Lioumis et al., 2012; Picht et al., 2009; Sondergaard et al., 2021; Tardelli et al., 2022; Weise et al., 2023). The neuronal responses to TMS can be measured through indirect brain activation, such as by observing peripheral muscle activation or behavioral outputs, and through direct brain activation with electroencephalography (EEG; Hernandez-Pavon et al., 2023; Ilmoniemi et al., 1997; Tremblay et al., 2019). TMS has been widely used to study the motor network due to the simplicity of measuring TMS-evoked motor evoked potentials (MEPs) with electromyography (EMG; Rossini et al., 2015). A number of commonly used stimulation protocols define the stimulation target and/or intensity with respect to (*i*) the motor hotspot (the cortical stimulation target that evokes the highest consistent motor response) and (*ii*) the resting motor threshold (rMT; the stimulation intensity needed to consistently evoke muscle responses). Thus, motor mapping is important, *e*.*g*., for many neuromodulation protocols (Caulfield et al., 2021; George et al., 2010; Groppa et al., 2012; Hoogendam et al., 2010; Lefaucheur et al., 2020; O’Reardon et al., 2007; Somaa et al., 2022; Veldema and Gharabaghi, 2022), and for locating muscle representational areas for surgical planning (Kantelhardt et al., 2010a; Krieg et al., 2017; Najib et al., 2011; Picht et al., 2009, 2016; Säisänen et al., 2010; Takahashi et al., 2013; Vitikainen et al., 2009). Reliable estimation of the hotspot and rMT requires high expertise and relatively long mapping times (∼10+ minutes; Harquel et al., 2017; Meincke et al., 2016; Tervo et al., 2020), and the result is dependent on the lab, equipment and the operator (Peterchev et al., 2012; Sondergaard et al., 2021). Furthermore, acquiring the expertise required is a slow process, introducing further errors and variability among new users. Therefore, automated, user-independent TMS targeting is needed for standardization, acceleration, and ensuring high precision of TMS mapping procedures across labs and studies.

When manually searching for the motor hotspot, a trained experimenter shifts and rotates the TMS coil intuitively around pre-determined regions to define the areas that produce the largest or most consistent MEPs. Manual mapping requires prior knowledge, such as the location of anatomical landmarks (e.g., hand motor knob; Yousry et al., 1997), and continuous recollection of the data observed so far to select where to target next. In algorithm development, this expert behavior needs mathematical formalization to create policies that intelligently select the next stimulation targets. Previous algorithms have utilized Bayesian optimization (Faghihpirayesh et al., 2021, 2020; Tervo et al., 2022, 2020), Bayesian modelling (Harquel et al., 2017), heuristic rules (Grab et al., 2018; Meincke et al., 2016), as well as grid approaches (Dormegny-Jeanjean et al., 2022), with varying results. The best precision was achieved by Meincke et al. with an average 1.4-mm precision in the estimated hotspot location, requiring over an hour of stimulation. The fastest convergence was reported by Tervo et al., (2020) with only an average of 18 samples needed to define the hotspot location with an accuracy of 1.4 mm and precision of 3.2 mm. However, in this case, the optimization was performed in just one dimension, along a straight line. A few of these studies have compared automated mapping with manual hotspot hunting (Dormegny-Jeanjean et al., 2022; Harquel et al., 2017; Meincke et al., 2016; Tervo et al., 2020). In these, automatic mapping indeed improved accuracy and/or precision. To our knowledge, however, no published study has yet demonstrated automatically finding both the optimal location and orientation simultaneously.

Fully automatic mapping requires machine-controlled shifting of the stimulation target, such as robotized TMS (for example: Grab et al., 2018; Matsuda et al., 2021; Noccaro et al., 2021; Pennimpede et al., 2013; Richter et al., 2013) or multi-locus TMS (mTMS; Koponen et al., 2018; Matsuda et al., 2024; Navarro de Lara et al., 2021; Nieminen et al., 2022; Sinisalo et al., 2024; Souza et al., 2022). Here, we utilize an mTMS system (Nieminen et al., 2022), which allows electronically shifting the stimulation target, such as stimulation location and orientation, without physically moving the coil set.

This study aims to find the motor hotspot as quickly and accurately as possible through a novel mapping algorithm, which is an extension of the original Bayesian Optimization Of Stimulation Targeting (BOOST) algorithm (Tervo et al., 2022, 2020). The novel 3D-BOOST algorithm simultaneously optimizes the location and orientation of the maximum electric-field (E-field) vector on the cortical surface. We demonstrate a fast and accurate hotspot mapping in the 3-dimensional search space by utilizing Bayesian optimization with Gaussian processes. The speed and accuracy of the developed algorithm was tested online by mapping the motor representation of the right first dorsal interosseus (FDI) muscle in 5 subjects with 3D-BOOST-driven mTMS.

## Methods

### Algorithm description

3D-BOOST is based on Bayesian optimization with Gaussian processes (Garnett, 2023a; Rasmussen and Williams, 2006). The log-transformed MEP peak-to-peak amplitude *y*_*n*_ is modelled as

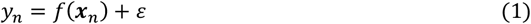

where *f*(***x***_*n*_) describes the log amplitude at target ***x***_*n*_ and *ε* is normally distributed noise with variance describing the variability of the MEP. Here, the dimensions of ***x***_*n*_ include two surface coordinates and an angle, describing the location and orientation of the stimulating E-field maximum. We assume the function *f*, describing the effect of changing the E-field (target ***x***_*n*_) on the recorded MEP amplitudes, to be smooth. Before any data collection, *f* is unknown. If we assume the values of *f*(***X***) with ***X*** = [***x***_1_, ***x***_2_,…, ***x***_*n*_] to be jointly Gaussian, we can write *f* in terms of a Gaussian Process (GP) model:

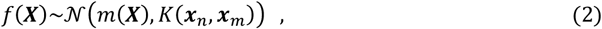

where *m*(***X***) and *K*(***x***_*n*_, ***x***_*m*_) denote the *a priori* expectation values of the mean of *f*(***X***) and the covariance between *f*(***x***_*n*_) and *f*(***x***_*m*_), respectively. Here, a squared exponential (SE) kernel is used for the covariance function, defined as:

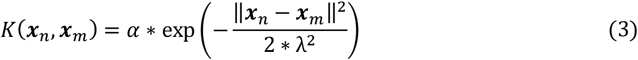

where the hyperparameters *α* and *λ* describe the variance of function values and the smoothness of the function *f*, respectively.

With these definitions, the posterior mean 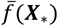 and variance cov(*f*(***X***_∗_)) functions, evaluated at targets ***X***_∗_, can be calculated by Bayesian inference given measurements ***y*** = [*y*_1_, *y*_2_,…, *y*_*n*_] using the following expressions:

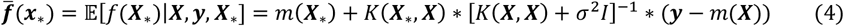

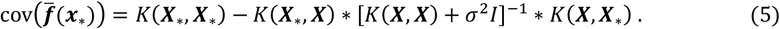

Here, the posterior mean describes the current model estimate of the average MEP size evoked from any potential stimulation target in ***X***_∗_ based on the MEP measurements ***y*** from already stimulated targets ***X***. The posterior variance describes the model uncertainty about the mean MEP size at each target ***X***_∗_.

To achieve a fast automatic optimization of the stimulation target such that the MEP response is maximized, the algorithm needs to choose the next sampling target intelligently. The aim is to find the stimulation target at which, when sampled, the algorithm gains maximum information about the optimization goal: the target yielding the maximum MEP response. The algorithm is guided to select sampling points that maximize the expected marginal gain in utility:

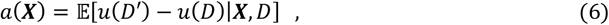

where *D′* is the dataset containing the new sample at ***x***_next_ . Defining the utility function *u* as 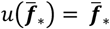, scaling linearly with the expected amplitudes of the response, the expected improvement (EI) acquisition function (Garnett, 2023b; Jones et al., 1998; Mockus, 1975) can be written as:

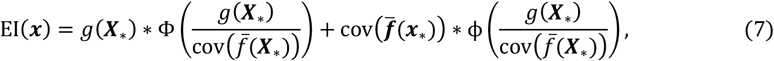

where

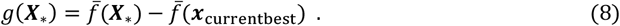

Φ and ϕ are the Gaussian cumulative distribution function and probability density function, respectively, and ***x***_currentbest_ is the current estimate of the best target. Intuitively, the EI acquisition function balances between sampling at targets with high uncertainty and sampling at targets with large posterior mean. This exploration–exploitation ratio can be adjusted by adding a small value *ϵ* to 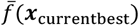 before calculating EI, where a larger *ϵ* shifts the behavior in a more explorative direction (Jones, 2001; Lizotte, 2009).

Finally, since the 3D-BOOST algorithm is designed to minimize the number of samples and since the marginal utility of additional samples diminishes over time, it is essential to establish a stopping criterion. Setting the criterion involves a compromise between the duration of data collection and accuracy: a lenient criterion halts sampling early, using less data at the expense of accuracy and precision in estimating the maximum, while a strict criterion ensures a more reliable estimate of the maximum but demands a larger number of samples and therefore increased experimental time. In this study, we tested stopping criteria utilizing the maximum deviation of the optimal target over the last *q* samples. Based on previous work and testing data, *q* was set to 15, and deviation thresholds to 1.5 mm and 5° after collecting at least 35 samples (Granö, 2023).

To define how the maximum evolves over time, the algorithm needs a definition of the best stimulation target. We used two definitions for this: (a) the GP posterior maximum location, that is, the target estimated to yield the largest MEPs according to the GP model, and (b) the center of gravity (CoG) of the measured MEPs.

The algorithm performs the following optimization loop:

1. Stimulate three to five pseudo-randomly chosen targets to initiate the dataset *D* and calculate the posterior mean and variance from the attained information (Eqs. 4, 5).
2. Based on the posterior mean and variance (current beliefs) of the MEP distribution, choose the next measurement/evaluation target ***x***_next_ by finding the maximum of the EI acquisition function (Eq. 7) .
3. Measure the MEP when stimulating ***x***_next_, append ***x***_next_ and the observed MEP amplitude *y*_new_ to *D* and update the posterior mean and variance (Eqs. 4, 5).
4. Repeat steps 2 and 3 until stopping criteria are met and return the best target to the user.

The algorithm contains three parameters that need to be set: the kernel variance scale *α*, the kernel length scale *λ*, and the EI exploration–exploitation parameter *ϵ* . Strategies for setting them were optimized on recorded test data (Granö, 2023). The variance scale *α* at each iteration was set as

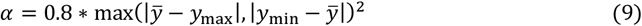

The length scale *λ* was set with the upcrossings formula (Rasmussen and Williams, 2006):

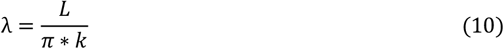

where *L* is the length of the search space, and *k* is the expected number of upcrossings. Based on running the algorithm on test data, *k* was set to 1.8. The parameter *ϵ* was set adaptively according to the measured responses (see Experiment Section for more details).

The implementation of the algorithm gives visual feedback to the user during the hotspot mapping. The algorithm’s graphical user interface is shown in Fig. 1.

**Figure 1.**
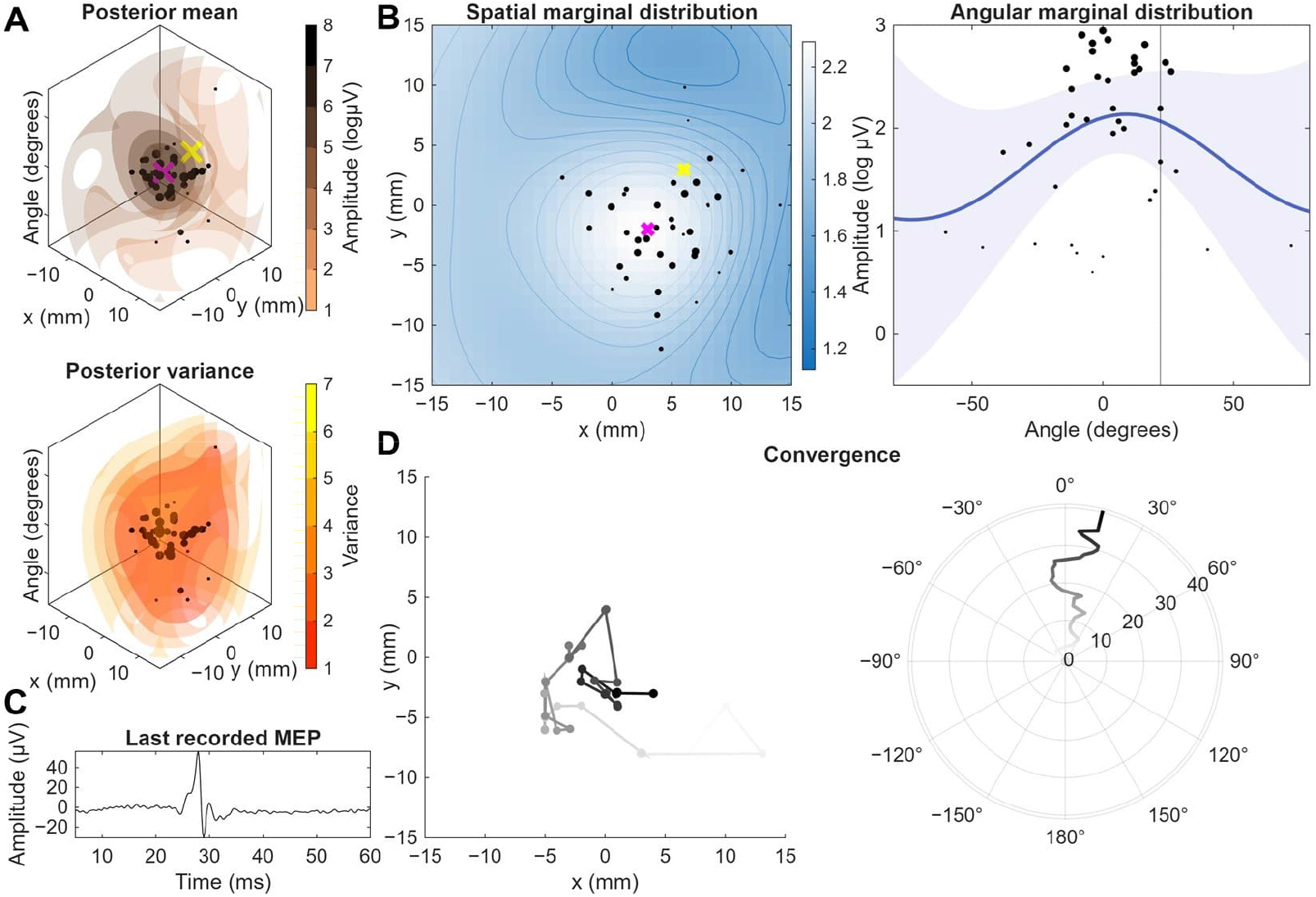
Graphical user interface. (A) The Gaussian process posterior mean and variance, showing the current estimate of the motor responses as a function of location and orientation. The black dots indicate the measured responses, and their sizes indicate the recorded amplitude. (B) The marginal spatial and angular distributions, visualized for clarification. In (A) and (B), the pink crosses indicate the current best estimate of the motor hotspot, and the yellow crosses (or vertical line in the angular marginal distribution) show the next sampling point. (C) The last recorded MEP response. (D) Traces showing how the estimate of the best target had evolved over the past samples, giving an indication of algorithm convergence. Left (location): Darker color indicates more recent estimates. Right (orientation): Orientation estimate as a function trial number (indicated by distance from origin).

To extract an MEP peak-to-peak amplitude, an epoch of –100…100 ms around the TMS pulse was extracted. A baseline correction was performed by subtracting the mean of the baseline calculated from –100 to –5 ms, after which a bandpass filter was applied with cutoff frequencies 5 and 1000 Hz. Then, a 50-Hz sine wave was fitted to the baseline time window, and the fit was extended to the whole window and subtracted from the signal to suppress 50 Hz line noise. If the signal range exceeded 60 µV in the baseline time period, the trial was repeated. The MEP peak-to-peak amplitudes were calculated from the EMG signal in the 10–60-ms time interval after the TMS pulse.

### Experiment: Real-time validation

A set of experiments was performed to characterize and validate the 3D-BOOST algorithm by running it online on five subjects (age range: 25–47 years, 1 left-handed, 3 male). All subjects signed a written informed consent, and the experiments were approved by The Coordinating Ethics Committee of the Hospital District of Helsinki and Uusimaa (HUS/1198/2016).

The subjects were seated in a comfortable chair and instructed to relax while keeping their eyes fixed on a cross three meters away. The subjects wore earplugs underneath earmuffs to protect their hearing. EMG was recorded with a TMS-compatible amplifier (NeurOne, Bittium Plc., Finland) from the FDI muscle, with the EMG electrodes placed in a bipolar belly–tendon montage. The ground electrode was placed on the styloid process of the ulna. The data were sampled at the sampling frequency of 5000-Hz in DC recording mode, lowpass filtered at 1250 Hz, and streamed to the computer that controlled the mTMS system.

TMS was delivered with a robot-controlled 5-coil mTMS system (Matsuda et al., 2024; Nieminen et al., 2022), allowing electronic adjustment of stimulation location and orientation within a cortical region of 3 centimetres in diameter. The mTMS coil set consisted of five overlapping coils, allowing the shape of the induced E-field to be modulated as a superposition of the fields induced by individual coils. The E-field strength was estimated based on a spherical 70-mm-radius volume conductor model 15 mm from the coil center. The coil currents for each stimulation target were optimized for each target and orientation (Laine, 2025) based on E-field distributions that had been measured for each coil with a search coil under the stimulator (Nieminen et al., 2015). The position, orientation, and tilt of the coil set were guided by the open-source InVesalius neuronavigation software (Amorim et al., 2015; Matsuda et al., 2025; Souza et al., 2018) with an infrared motion capture setup consisting of eight cameras (OptiTrack; NaturalPoint Inc., USA) and automated robotic control with 1-mm and 3° precision to keep the coil set at the target location (Matsuda et al., 2021; Elfin Collaborative Robot, Shenzhen Han’s Robot Co., Ltd., China). Neuronavigation was based on individual T1-weighted magnetic resonance images (MRIs).

First, the motor hotspot and resting motor threshold (rMT) of the FDI muscle in the right hand were manually determined. The motor hotspot was defined as the cortical region producing the largest MEP amplitudes in the target muscle, and the rMT as the stimulation E-field intensity eliciting MEPs with peak-to-peak amplitudes larger than 50 µV in 10 out of 20 repetitions (Rossi et al., 2021) when stimulating in the optimal location and orientation. The coil set was placed over the primary motor cortex (M1) and the hand motor knob was mapped by electronically shifting the stimulation target until a target yielding consistent MEPs was found. If no clear hotspot was found, the coil set was robotically moved, after which the electronic search was repeated. Defined rMTs were (mean ± SD) 91 ± 7 V/m, range 84–100 V/m.

After the motor hotspot and threshold were found, 150 stimuli were delivered around the hotspot at 110% rMT to validate that the true hotspot was well defined within the search space of the algorithm. This mapping was performed by electronically shifting the E-field maximum while keeping the position of the coil set fixed with respect to the head. The inter-stimulus interval (ISI) was drawn pseudo-randomly between 3 and 3.5 s. However, at times, the mTMS coil channels may require more time to charge or discharge, in which case ISI could extend up to 5 s. Stimulation was only performed if the coil set was within 1 mm and 3° from the coil target. The stimuli were arranged in a grid with a higher density of grid points near the manually found motor hotspot.

Next, 3D-BOOST was run three times with each of the three tested sampling policies (explorative, intermediate, and exploitative; Table 1) with a randomized run order and ISI as described above, totaling nine runs per subject. For each run, 100 pulses were delivered at 110% rMT. The search space was circular with 15-mm radius and 1-mm spacing between grid points and with –80 to 80° range with 1° spacing in the angular dimension. The 0° orientation was defined as the direction of the coil, set approximately perpendicular to the central sulcus. At the end of the experiment, rMT was remeasured at the globally best spot, defined as the CoG calculated from all recorded data.

**Table 1.**
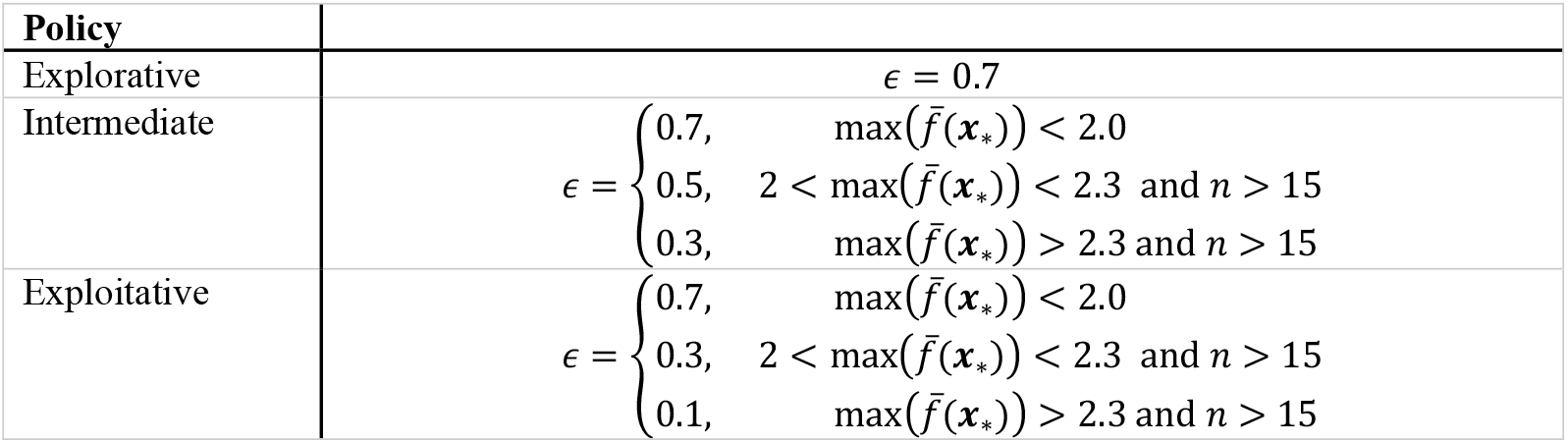
The three different parameter sets for the exploration/exploitation parameter *ϵ* . The number of samples recorded in the run is denoted by *n*. The parameters were defined based on the performance of the algorithm on test data.

### Data analysis

The performance of the algorithm was estimated with two metrics. The first metric was accuracy, calculated as the distance from the recommended target of the individual runs to the ground truth best spot. The ground truth best spot was defined as the CoG calculated from all data of each subject. For each such comparison, the global CoG was calculated, excluding the MEPs recorded during that run. The second metric was the expected MEP amplitudes at each target recommended by the individual runs. We calculated the posterior mean from all recorded MEPs (appr. 1050 in total) from the given subject to evaluate a ground truth prediction for MEP amplitudes everywhere in the search space. From this model, we predicted the MEP amplitudes at the stimulation targets suggested by individual algorithm runs. We assessed the optimality of those targets by computing the ratio between the resulting MEP amplitudes and the ground truth maximum MEP amplitude (prediction at the ground-truth GP maximum).

The algorithm-found best targets were defined in two different ways, as the CoG and the GP maximum location, calculated by utilizing the samples taken during the run. The accuracy of the algorithm was further assessed with post-hoc early stopping; namely, only the samples taken until fulfillment of the stopping criteria were considered for defining the algorithm-found optimal targets. We used a linear mixed-effect model (LMM) with restricted maximum likelihood estimation to assess the following fixed effects on accuracy: (1) sampling policies, (2) definition of best target (GP maximum and CoG), (3) convergence criteria, and (4) subject-specific stimulation intensity. The subject-specific intensity was calculated with respect to the rMT defined at the end of the experiment at the global CoG, to take into account that the initial rMT might have been estimated at a suboptimal spot. A random effect was included for each subject. We checked the normality and homoscedasticity of the residuals with quantile–quantile plots and standard vs. fitted value visualizations. We then performed an analysis of variance (ANOVA) on the fitted model by computing F-tests for each fixed effect utilizing Satterthwaite’s approximation. For each model coefficient, we estimated the number of subjects needed to reach significance using the model-derived fixed-effects estimates and standard errors (SE), assuming that SE scales with 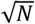, where *N* is the number of subjects (see Supplementary material).

The average accuracy of the algorithm-recommended targets was further compared with the accuracy of manually chosen target with a separate LMM, with a single fixed effect (manual vs. all algorithm-defined targets pooled over conditions) and subject-specific random intercepts.

Separate LMMs were computed for the location and orientation. The significance threshold was set to *p =* 0.05 for all tests, and multiple comparisons were Bonferroni-corrected.

## Results

### Motor maps

In all subjects, the global CoG was well defined within the search space of the algorithm (distance from the coil-set center: mean ± SD 4.2 ± 2 mm, range 2.1–7.9 mm), indicating that the search space sufficiently covered the relevant brain region. Manual motor mapping took 28 ± 13 minutes, and the manually defined hotspot was (mean ± 95% CI) 4.0 ± 2.8 mm and 11 ± 7° from the global CoG. The rMT was identical or lower at the end of the experiment when measured from the global best target than the rMT defined at the beginning of the experiment (mean difference ± SD: −4 ± 4 %) in all except one subject, in which the rMT increased by 12%. The acquired motor map with all recorded data from one subject is shown in Fig. 2.

**Figure 2.**
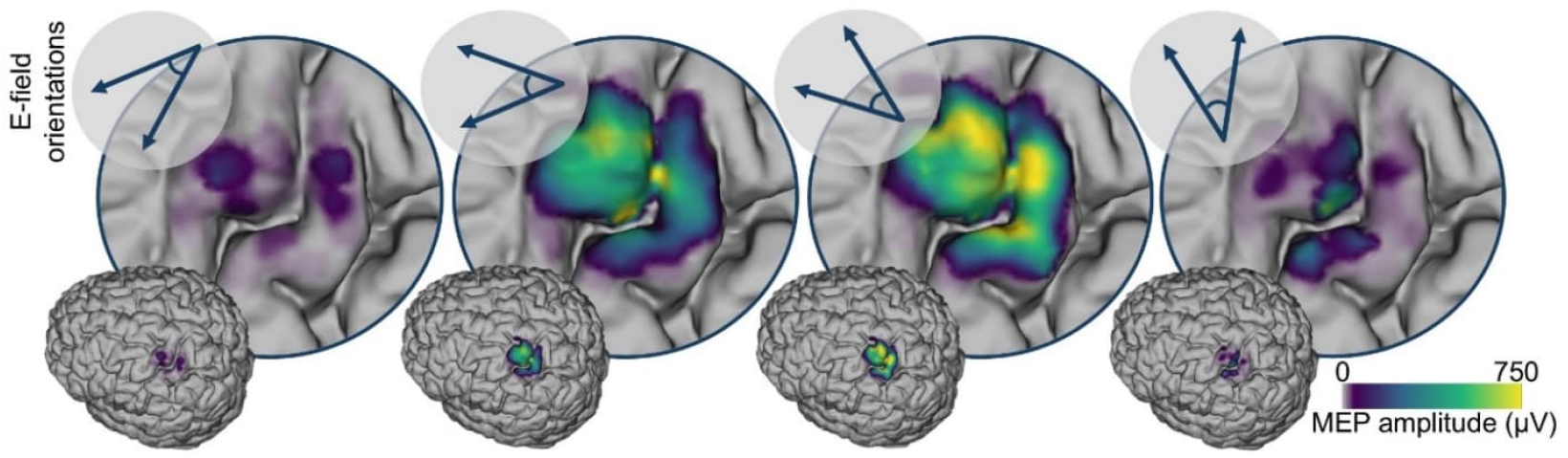
Motor maps generated by projecting the measured MEPs to the cortical surface and applying Gaussian smoothing in a representative subject. From left to right, the stimulus orientation changes with 40° steps. The maps were generated from a total of 1045 trials.

### Algorithm performance

3D-BOOST was able to find stimulation targets close to the global CoG across all runs. The distances from the algorithm-found targets to the global CoG, both when accounting for the full 100 samples taken during each run as well as with early stopping are shown in Fig. 3. Out of all the conditions, the most explorative condition (condition 1) with the full 100 samples and CoG as the definition of the optimal target performed the best, with an average distance (± 95% CI) to the global CoG of 1.7 ± 0.5 mm and 4 ± 2°. With early stopping at convergence, the average distance increased to 2.1 ± 0.7 mm and 6 ± 2°, while requiring only about half of the samples (47 ± 6 samples). The mean, 95% CI, and standard deviations of the distances from the algorithm-recommended targets to the global CoGs for each condition are reported in the Supplementary material.

**Figure 3.**
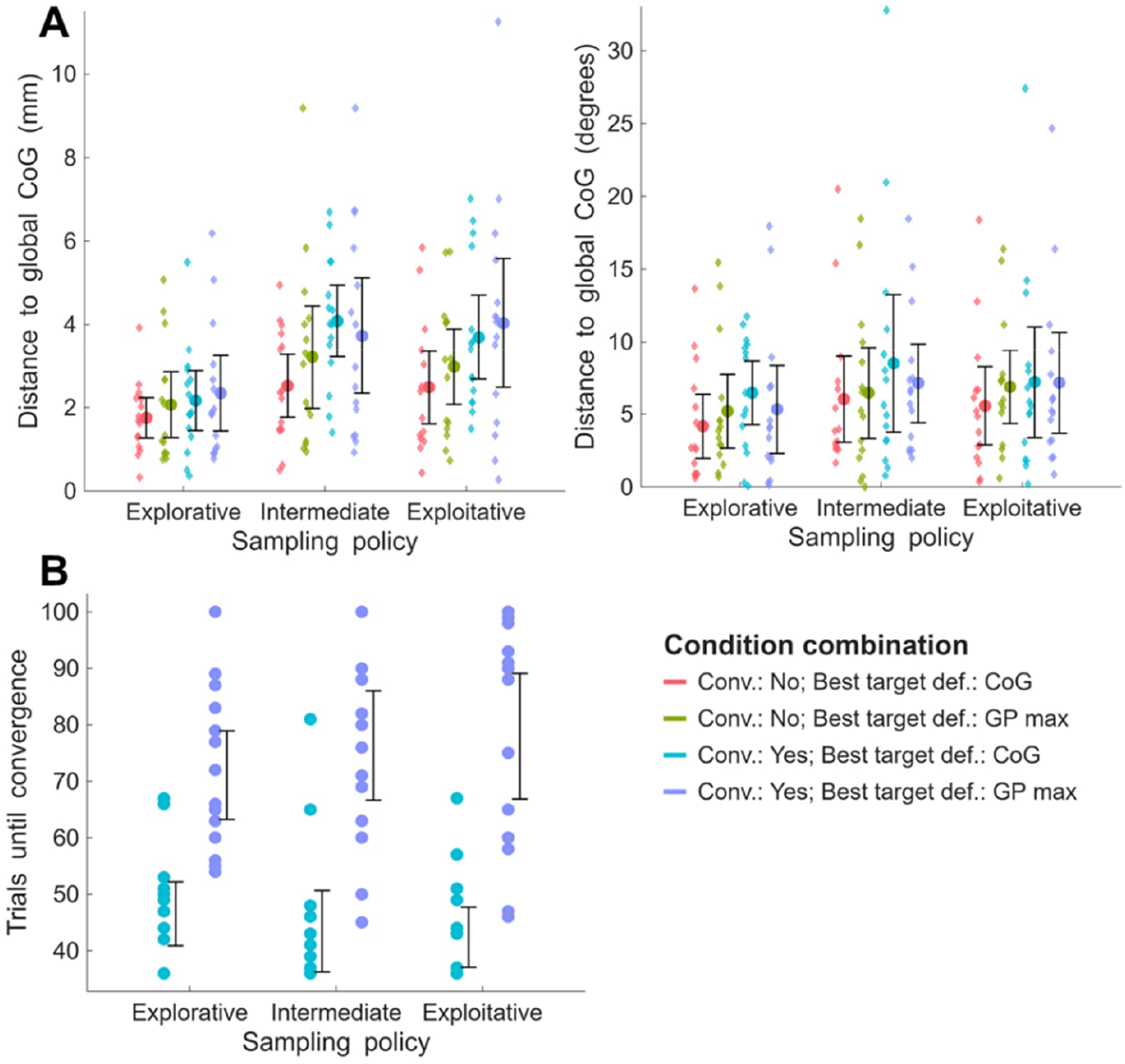
(A) The average distances from the algorithm-recommended targets to the global centers of gravity (CoGs) for each condition (sampling policy, definition of best target, and convergence criteria) are shown as colored dots, with 95% confidence intervals (CIs; error bars), and distances of individual runs are presented with small diamonds. (B) The number of trials required until fulfillment of the convergence criteria for each sampling policy and both optimal target definitions (CoG vs Gaussian process maximum). When the CoG was defined as the optimal target, convergence was reached faster.

In general, defining the best parameters with the CoG yielded a smaller error than the GP maximum. The GP maximum resulted in smaller error than the CoG only when utilizing a convergence criterion with the intermediate sampling policy (CoG: 4.1 mm and 9°; GP Max: 3.7 mm and 7°), but requiring approximately 30 samples more to do so. Utilizing the full 100 samples resulted in a higher accuracy than when applying a stopping criterion. The algorithm-found targets for all subjects, as well as the data from one representative subject, are presented in Fig. 4.

**Figure 4.**
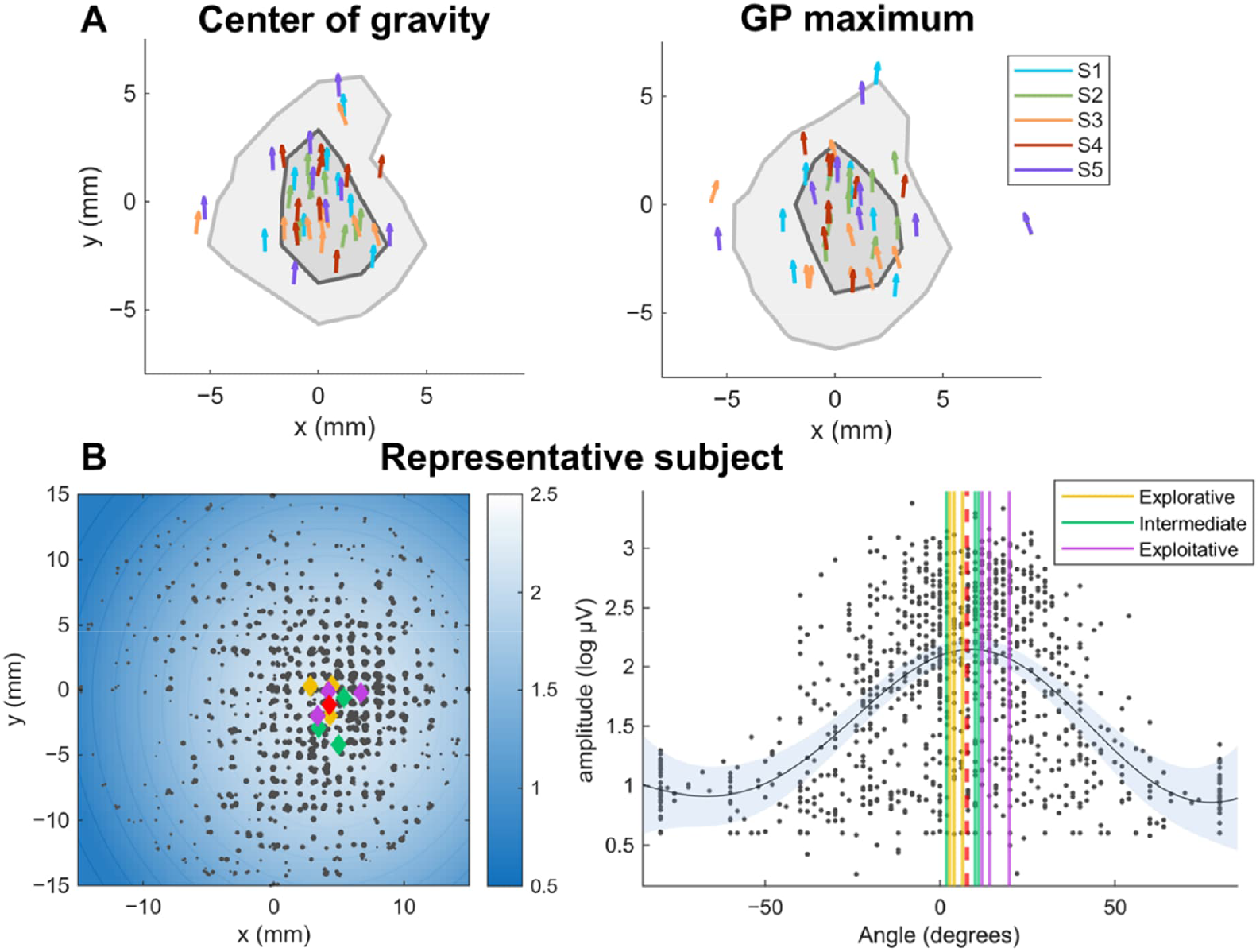
(A) The algorithm-found optimal targets for all the runs for all subjects (colored arrows, where the direction indicates the target orientation) for both best target definitions (center of gravity (CoG) vs Gaussian Process (GP) maximum), utilizing the full set of 100 samples per run. For each subject, the origin of the coordinate system is set to the subject-specific global CoG and 0° (upward) as the optimal orientation. In half of the runs, a target was found within the darker shaded region, and 95% of the time within the lighter shaded region. (B) All recorded motor-evoked potentials (MEPs) for one representative subject, with the dot size indicating the MEP size. The gradient in the left figure shows the spatial marginal distribution of the GP posterior mean, and the black curved line in the right figure represents the angular marginal distribution. The blue shaded area shows the model uncertainty. The red diamond (left figure) and dashed vertical line (right figure) show the global CoG for this subject, and the other colored diamonds and vertical lines indicate the algorithm-found best targets.

The expected MEP amplitudes from the algorithm-found optimal stimulation targets were similar to the expected MEP amplitude at the GP posterior maximum calculated on all recorded data. The most explorative policy with the CoG yielded 99% of the maximum MEP amplitudes on average. Table 2 lists all expected MEP amplitudes of the various conditions as a percentage of maximum expected MEP amplitudes.

**Table 2.**
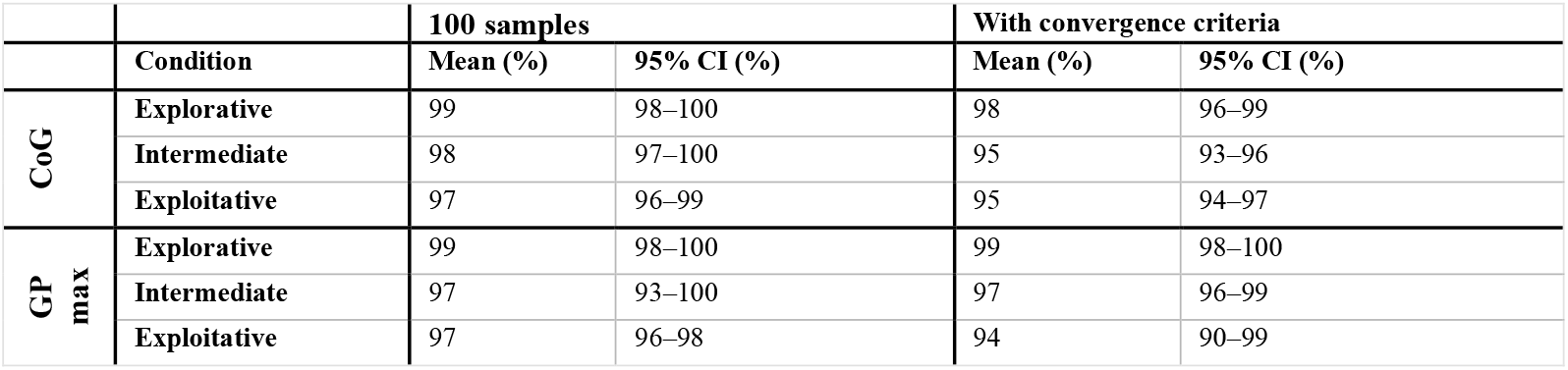
The expected mean MEP amplitude of the algorithm-found targets, as a percentage of the maximum MEP amplitude. The MEP amplitudes for each target were evaluated from the GP posterior calculated from all recorded MEPs (appr. 1050 in total) from the given subject. The values are reported for all conditions (sampling policies, definition of best target, and convergence criteria).

The algorithm-found targets were in general closer to the global CoG with weaker stimulation intensities, apart from the GP maximum condition utilizing convergence criteria. This dependency on the stimulation intensity appeared more prominent with more exploitative sampling paradigms (Fig. 5). A similar pattern was observable across the found stimulation angles, with the exception of small differences across sampling policies in the subject for whom the highest stimulation intensity was used. However, the LMM did not signify that the stimulation intensity played a significant role (*p* = 0.24).

**Figure 5.**
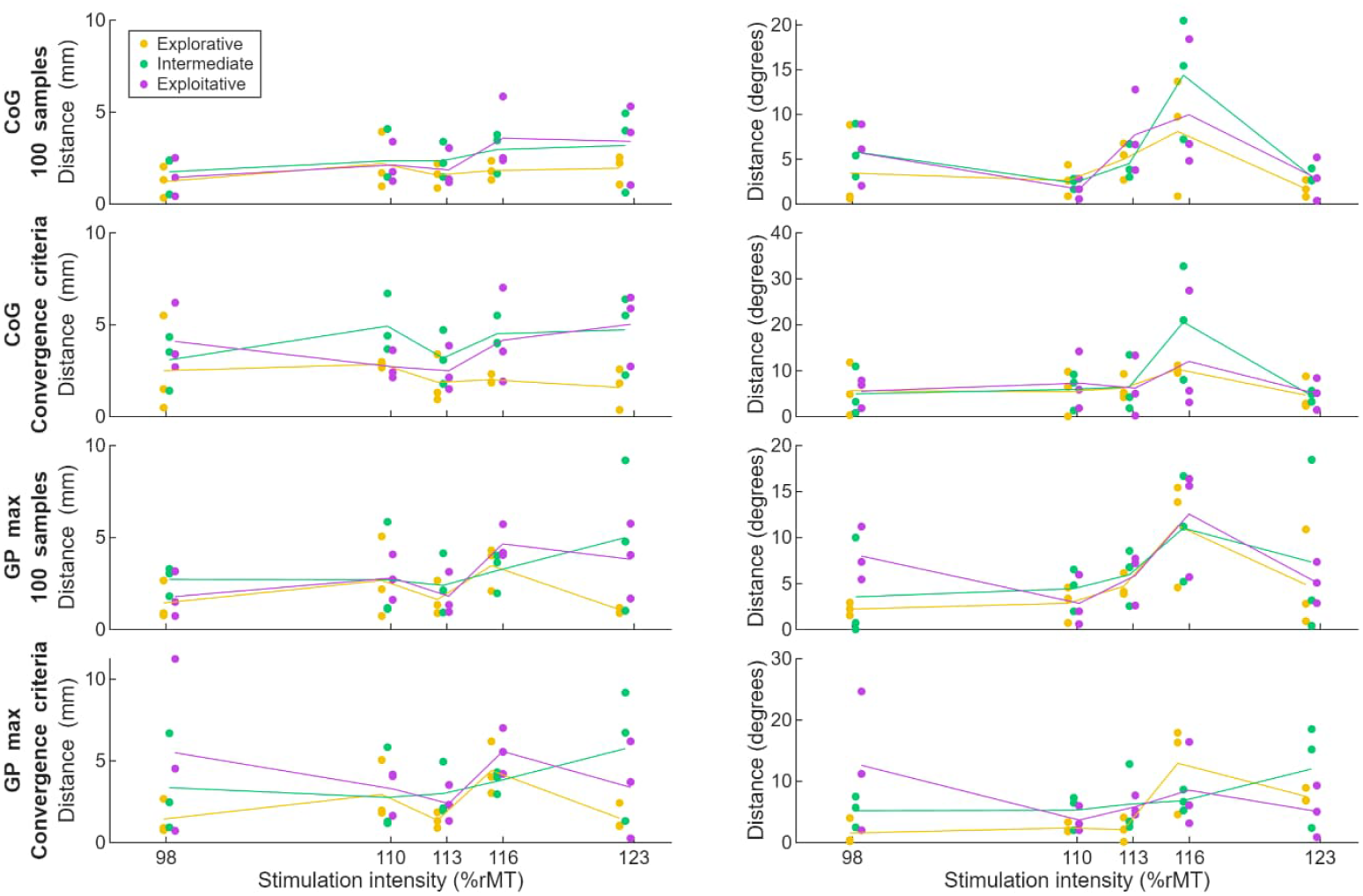
The distances from the algorithm-found optimal targets to the global CoG. The distance appeared to grow with higher stimulation intensity with more exploitative sampling paradigms, as observed across subjects. The lines show the average distance from the global CoG with respect to the rMT at the end of the experiment. The LMM did not suggest this trend to be significant.

With the most explorative sampling policy, increasing the number of samples decreased the distance to the global CoG the fastest (Fig. 6), which is accentuated when defining the optimal target as the GP maximum. This is also reflected in the convergence rates with the GP maximum; the other sampling policies often did not fulfil the convergence criteria even with 100 samples. When using the CoG, the more exploitative policies often converged with the minimum number of samples (mode: 37 samples), while the most explorative policy tended to sample slightly more before fulfilling the convergence criterion (mode: 48 samples).

**Figure 6.**
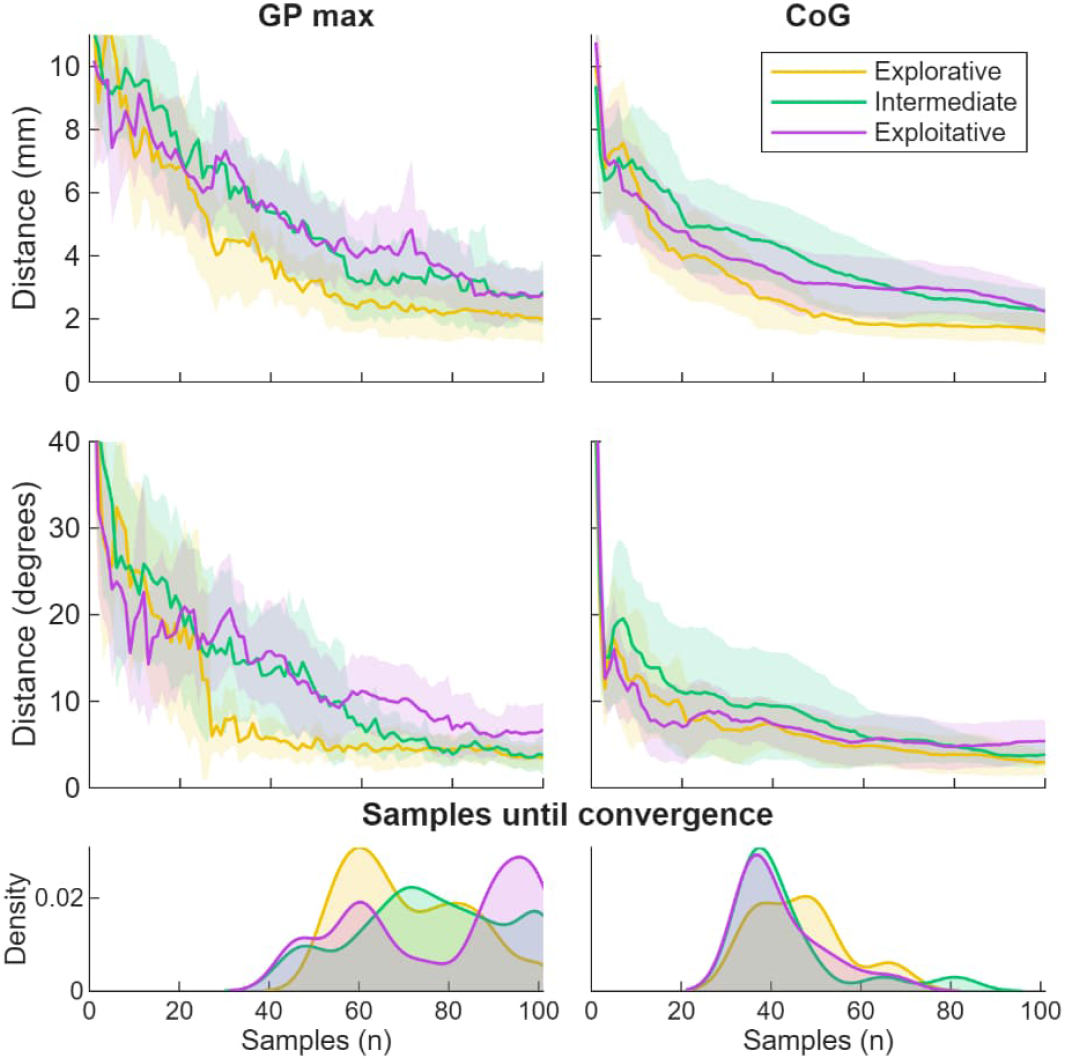
The average distance to the global CoG with increasing samples, for all conditions (CoG vs. GP as the best target definition; sampling policy). The shaded regions indicate the 95% confidence intervals in the top and middle rows. The bottom row shows the number of samples required until the convergence criterion was met for the different conditions, estimated as a probability density.

The LMM did not suggest any differences in algorithm performance across any of the conditions. The results for all the model parameters and post-hoc power analysis results are reported in the Supplementary material. There was no significant difference between the manual and algorithm-defined target locations according to the LMM (location: *p* = 0.19, orientation: *p* = 0.057).

## Discussion

We developed an algorithm, called 3D-BOOST, capable of automatically identifying the motor hotspot location and orientation without physical movement of the stimulation coils. We demonstrated fast and accurate mapping with mTMS and Bayesian optimization, with an average estimated error of 1.7 mm and 4° for the best performing condition with 100 TMS pulses, and 2.1 mm and 6° when applying a convergence criterion and utilizing only 47 trials on average. This is a significant development from the original BOOST algorithm (Tervo et al., 2020), which optimized the E-field in only one dimension (orientation or translation along a line) with an accuracy of 1.4 mm or 3.2°, respectively. Optimization in three dimensions that only requires slightly more than double the samples than in one dimension is an important step toward fully automated mapping approaches. Furthermore, 3D-BOOST is an improvement over other automatic mapping approaches reported. For instance, while better precision of 1.4 mm was reported by Meincke et al. (2016), it required one hour of stimulation, rendering the method impractical in clinical settings. Harquel et al. (2017) reported a precision of 8.2 ± 5.4 mm and Dormegny-Jeanjean et al. (2022) 7.7 ± 8 mm, with somewhat comparable numbers of stimuli to those in this work (13.5 ± 3.3 locations with 3–5 samples each, and 100 samples, respectively). Faghihpirayesh et al. (2021, 2020) required 122.4 ± 27.5 and 150–282 samples, respectively, but reported neither accuracy nor precision, instead examining model prediction and MEP size agreement.

Importantly, 3D-BOOST reduced the measurement error by half when compared to the manual mapping (2.1 mm and 6° vs. 4.0 mm and 11°). A few previous studies have reported manual mapping errors of 3.5 mm (with only linear translation; Tervo et al., 2020), 4.4 mm (calculated as mean range between CoGs; Zdunczyk et al., 2013), 4.8 mm (when compared to direct electrical stimulation of the cortex; Paiva et al., 2024), and 19 mm (reported as distance between active and passive motor mappings; Dormegny-Jeanjean et al., 2022). Although these numbers are not directly comparable due to differences in methodology, 3D-BOOST appears to achieve a higher accuracy than that of a human operator.

It is important to note that the “best target” for evoking MEPs is not uniquely defined, as MEPs are evoked from a large area. In this work, both the GP posterior maximum, i.e., the target yielding the largest MEPs according to the GP model, and the CoG were compared. We observed that the GP maximum is more likely to be disproportionately affected by small numbers of occasional large deviations in MEP response amplitudes, rendering it less reliable and repeatable than the CoG, which is a well-established method for determining the motor hotspot (for example; Jin et al., 2023, 2022; Malcolm et al., 2006; Nazarova et al., 2021; Rossini et al., 2015; Sondergaard et al., 2021; Tardelli et al., 2022). The CoG is more robust, but can be biased towards the center of the search space. This could be particularly problematic in cases of poor coil set placement, where the algorithm is unable to sample evenly from the whole MEP distribution. Even though we defined the best target for algorithm validation, the sampling process does not require an estimate of the optimal target in order to choose the next sampling point. Furthermore, the implemented UI of 3D-BOOST visualizes the whole mapped distribution, ensuring that the operator remains aware of the full motor map and mapping procedure.

The statistical analysis did not suggest significant effects of any of the conditions on the algorithm performance. However, as we report the results with only five subjects, the power of the statistical analysis is limited. These results should therefore be regarded as preliminary. According to post-hoc power analysis (see details in Supplementary material), 13 subjects would have been needed to observe a statistically significant effect of the sampling policy choice, 12 for a significant effect of the choice of using convergence criteria, and 59 for a significant effect of the definition of the best target (CoG vs. GP maximum). However, the current preliminary results suggest that any potential effect of the specific parameter choices on the overall algorithm performance are only small to moderate. The data collection is still ongoing, and future versions of this work will include an expanded dataset to enable more robust statistical testing.

The current version of the 3D-BOOST algorithm does not yet automatically select or find the stimulation intensity for the mapping procedure. Therefore, it is possible for the user to select a suboptimal intensity. While using an insufficient intensity could yield only a few MEPs, resulting in an unreliable mapping, an unnecessarily high intensity could produce a saturated MEP distribution with poor specificity. In this work, we aimed to run the algorithm with 110% rMT; however, due to small inaccuracies in our initial hotspot estimate and changes in corticospinal excitability over the course of the experiment, the actual mappings were performed at both slightly lower and higher intensities, confirmed by calculating the intensities used relative to the rMT at the end of the experiment (range: 98 to 122% rMT). Our post-hoc visualizations suggested that the intensity played a larger role with the more exploitative paradigms, reflecting the higher risk of excessively sampling regions with large MEPs. However, more subjects are needed to confirm these observations. For practical use of 3D-BOOST, we recommend observing a few MEPs in the search space before starting the automatic mapping to ensure a sufficient stimulation intensity.

The algorithm chooses the first stimulation targets randomly; however, to further improve the performance and usability of automated hotspot hunting algorithms, anatomical priors for coil placement and initial search areas could be advantageous. Currently, the algorithm relies on the user to place the mTMS coil set in an appropriate position, introducing a risk of poor coil set positioning, which might affect the algorithm performance. Such priors might not only be advantageous for 3D-BOOST, but also for manual and robotic motor mapping procedures.

Furthermore, with better E-field modelling, the stimulated spots on the cortical surface could be determined more accurately. Currently, 3D-BOOST provides the user with the optimal E-field shift which maximizes the MEP response. This shift is given in millimeters and degrees, approximating moving the E-field maximum on a spherical surface by that amount. However, these units might not exactly correspond to the location of the cortical hotspot (Weise et al., 2023), even after having found the E-field that stimulates the cortical hotspot more strongly than other available E-fields. More accurate estimation of the cortical hotspot location requires realistic E-field modelling, which has recently been incorporated and demonstrated with mTMS in real time (Soto et al., 2025, 2023). Future version of the algorithm could utilize the capabilities of such modelling, allowing better anatomically informed mappings.

Using mTMS, the 3D-BOOST mapping process took approximately 3.6 minutes to reach convergence on average, but this mapping time was affected by the long capacitor charging/discharging delays between pulses. However, the mapping could be further accelerated by utilizing pulse-width-modulation (Sinisalo et al., 2025), which eliminates the need for large changes in capacitor voltages between pulses. With this method, ISI could be reduced to 1 second or shorter, which has been suggested to not significantly change the acquired motor maps (Mathias et al., 2014; van de Ruit et al., 2015). Thus, an mTMS system utilizing both pulse-width-modulation and 3D-BOOST could potentially map a muscle-representation area in under one minute.

Although the present 3D-BOOST algorithm is designed for the mTMS system, it can be adapted for robot-guided or manually performed conventional TMS; however, additional time would be required for physical coil adjustments compared to the rapid electronic shifts enabled by the mTMS technology (Sinisalo et al., 2024). The logic of 3D-BOOST can be applied to conventional TMS by translating mTMS-specific stimulation commands into commands for a robot moving a conventional coil, or visual guidance for a human operator. Estimating that physically moving the TMS coil with high precision by hand would take 8 s and by robot 2 s (Kantelhardt et al., 2010b; Richter et al., 2010), the required mapping time translates to approximately 6 and 1.5 min, respectively, using the average of 47 stimuli achieved by 3D-BOOST. As motor mapping is reported to usually take approximately 10 minutes or more (Harquel et al., 2017; Meincke et al., 2016; Tervo et al., 2020), a mapping time of 6 minutes or less would result in a moderate to significant acceleration also without the fast electronic targeting enabled by mTMS.

3D-BOOST can theoretically be adapted to any type of brain responses. The feasibility of 1D-optimization of stimulation orientation based on EEG responses was already demonstrated in Tervo et al., (2022). Recently, optimization of stimulation parameters (orientation, location, and intensity) was demonstrated by Parmigiani et al. (2025), further underlining the feasibility and utility of automatic mapping approaches also for TMS–EEG data. Their algorithm was designed to specifically increase early TEP reliability in the dorsolateral prefrontal cortex, which is a challenging target in TMS–EEG due to prevalent muscle artefact contamination (Mutanen et al., 2013). However, their approach may not directly generalize to other purposes, whereas 3D-BOOST can flexibly accept any feature for optimization. For example, maximizing the effective connectivity between two cortical regions may be of interest for indirect stimulation of areas challenging to target directly with TMS. In addition to EEG-based metrics, Bayesian optimization has been demonstrated to successfully improve stimulation parameter tuning in neurostimulation modalities such as epidural spinal cord stimulation for motor rehabilitation (Zhao et al., 2021) and invasive cortical stimulation for treatment of chronic post-stroke pain (Dastin-van Rijn et al., 2021).

By automating TMS mapping, we can standardize targeting across users and laboratories, bring personalized approaches to the clinics with accelerated and user-agnostic strategies, and improve targeting accuracy. This standardization has the potential to improve clinical outcomes and reproducibility of results and lessen the burden of training new operators.

## Conclusions

We demonstrated automatic, fast, and accurate TMS motor mapping with Bayesian optimization combined with multi-locus TMS, achieving 2.1-mm and 6° mean accuracy with as few as 47 stimuli on average. This is a significant improvement from previous algorithms in terms of speed and accuracy for finding an optimal stimulation target and orientation. This paves the way for faster experimental processes and more streamlined clinical applications, reducing costs, improving reproducibility and increasing the clinical utility of TMS.

