## Supplementary material for "Fast and standardized motor-hotspot determination with automated TMS mapping"

#### Algorithm performance metric: distance to global center of gravity

The algorithm’s performance was estimated by calculating the distance from the recommended target from the individual runs to the ground truth best spot. The ground truth best spot was defined as the center of gravity (CoG) calculated from all recorded data from each subject. For each such comparison, the global CoG was calculated excluding the MEPs recorded during that run. The algorithm-found best targets were defined in two different ways, as the CoG and the GP maximum location, calculated by utilizing the samples taken during the run. The algorithm’s accuracy was further assessed with post-hoc early stopping; namely, only the samples taken until fulfillment of the stopping criteria were considered for defining the algorithm-found optimal parameters. The mean ± 95% confidence intervals (CIs) and standard deviations (SDs) are reported in Supplementary table 1.

**Supplementary table 1.** The average distance, 95% confidence intervals (CIs) and their standard deviations from the algorithm-recommended targets to the global CoG, for each condition (sampling policy, definition of best target, and convergence criteria).

| | | 100 samples | | With convergence criteria | | |
|---|---|---|---|---|---|---|
| | **Policy** | **Mean ± 95% CI (mm; °)** | **SD** | **Mean ± 95% CI (mm; °)** | **SD** | **Convergence (mean±95%CI) (samples)** |
| **CoG** | **Explorative** | 1.7 ± 0.5; 4 ± 2 | 0.9; 4 | 2.1 ± 0.7; 6 ± 2 | 1.3; 6 | 47±6 |
| | **Intermediate** | 2.5 ± 0.7; 6 ± 3 | 1.3; 6 | 4.1 ± 0.9; 9 ± 5 | 1.6; 9 | 43±8 |
| | **Exploitative** | 2.5 ± 0.9; 6 ± 3 | 1.6; 6 | 3.7 ± 1.0; 7 ± 4 | 1.8; 7 | 42±6 |
| **GP max** | **Explorative** | 2.1 ± 0.8; 5 ± 3 | 1.4; 5 | 2.3 ± 0.9; 5 ± 3 | 1.6; 5 | 71±8 |
| | **Intermediate** | 3.2 ± 1.2; 6 ± 3 | 2.2; 6 | 3.7 ± 1.4; 7 | 2.5; 7 | 76±10 |
| | **Exploitative** | 3.0 ± 0.9; 7 ± 3 | 1.6; 7 | 4.0 ± 1.5; 7 ± 3 | 2.8; 7 | 78±12 |

#### Linear mixed-effects modelling and post-hoc power analysis

We applied a linear mixed-effect model (LMM) with restricted maximum likelihood estimation to assess the following effect on algorithm accuracy: (1) different sampling policies, (2) definition of best target (GP maximum and CoG), (3) use of convergence criteria, and (4) subject-specific stimulation intensity. The subject-specific intensity was calculated with respect to the rMT defined at the end of the experiment at the global CoG. A random effect was included for each subject. We checked the normality and homoscedasticity of the residuals with Q-Q plots and standard vs. fitted value visualizations. We then performed an analysis of variance (ANOVA) on the fitted model by computing the F-tests for each fixed effect utilizing Satterthwaite's approximation. For each model coefficient, we estimated the number of subjects needed to reach significance using the model-derived fixed-effects estimates and standard errors (SE), and assuming that the SE scales with $\sqrt{N}$, where $N$ is the number of subjects.

The average accuracy of the algorithm-recommended targets was further compared to the accuracy of manually chosen target with a separate LMM, with a single fixed effect (manual vs. all algorithm-defined targets pooled over conditions) and subject-specific random intercepts.

Separate LMMs were computed for the location and orientation. The significance threshold was set to $p$ = 0.05 for all tests and multiple comparisons corrected with a Bonferroni correction.

### LMM 1: Assessing the effect of conditions on location error

**Supplementary Table 2.** The F-statistics, *p*-values and estimated degrees of freedom for each fixed effect, calculated with ANOVA on the linear mixed-effects model output with Satterthwaite's method. The LMM was modelling the location error term.

| Term | F-statistic | *p*-value | Degrees of freedom (numerator, denominator) |
|---|---|---|---|
| Sampling policy | 0.95 | 0.39 | (2, 167) |
| Definition of best target | 0.25 | 0.62 | (1, 167) |
| Convergence | 0.43 | 0.51 | (1, 167) |
| Stimulation intensity | 1.4 | 0.24 | (1, 167) |
| Sampling policy × Definition of best target | 0.087 | 0.92 | (2, 167) |
| Sampling policy × Convergence | 1.87 | 0.42 | (2, 167) |
| Definition of best target × Convergence | 0.024 | 0.88 | (1, 167) |
| Sampling policy × Definition of best target × Convergence | 0.33 | 0.72 | (2, 167) |

**Supplementary Table 3.** The estimated number of subjects needed to reach significance for each model coefficient, calculated from the model-derived fixed-effects estimates and standard errors (SE), and assuming that the SE scales with $\sqrt{N}$, where $N$ is the number of subjects.

| Model coefficient | Subjects |
|---|---|
| Sampling policy: Intermediate | 13 |
| Sampling policy: Exploitative | 15 |
| Definition of best target: GP maximum | 78 |
| Convergence: Full | 46 |
| Stimulation intensity | 14 |
| Sampling policy: Intermediate × Definition of best target: GP maximum | 111 |
| Sampling policy: Exploitative × Definition of best target: GP maximum | 467 |
| Sampling policy: Intermediate × Convergence: Full | 12 |
| Sampling policy: Exploitative × Convergence: Full | 24 |
| Definition of best target: GP maximum × Convergence: Full | 801 |
| Sampling policy: Intermediate × Definition of best target: GP maximum × Convergence: Full | 38 |
| Sampling policy: Exploitative × Definition of best target: GP maximum × Convergence: Full | 50826 |

The LMM for modelling the difference between algorithm-found and manually defined targets returned an F-statistic of 1.74 and *p*-value of 0.19, with (numerator, denominator) (1, 183) degrees of freedom.

### LMM 1: Assessing the effect of conditions on orientation error

**Supplementary Table 4.** The F-statistics, *p*-values and estimated degrees of freedom for each fixed effect, calculated with ANOVA on the linear mixed-effects model output with Satterthwaite's method. The LMM The LMEM was modelling the orientation error term.

| **Term** | **F-statistic** | ***p*-value** | **Degrees of freedom (numerator, denominator)** |
|---|---|---|---|
| Sampling policy | 0.59 | **0.56** | (2, 167) |
| Definition of best target | 0.33 | 0.57 | (1, 167) |
| Convergence | 1.6 | 0.20 | (1, 167) |
| Stimulation intensity | 0.20 | 0.66 | (1, 167) |
| Sampling policy × Definition of best target | 0.061 | 0.94 | (2, 167) |
| Sampling policy × Convergence | 0.061 | 0.94 | (2, 167) |
| Definition of best target × Convergence | 0.73 | 0.39 | (1, 167) |
| Sampling policy × Definition of best target × Convergence | 0.026 | 0.97 | (2, 167) |

**Supplementary Table 5.** The estimated number of subjects needed to reach significance for each model coefficient, calculated from the model-derived fixed-effects estimates and standard errors (SE), and assuming that the SE scales with $\sqrt{N}$, where $N$ is the number of subjects.

| **Model coefficient** | **Subjects** |
|---|---|
| Sampling policy: Intermediate | 18 |
| Sampling policy: Exploitative | 32 |
| Definition of best target: GP maximum | 59 |
| Convergence: Full | 12 |
| Stimulation intensity | 97 |
| Sampling policy: Intermediate × Definition of best target: GP maximum | 359 |
| Sampling policy: Exploitative × Definition of best target: GP maximum | 1553 |
| Sampling policy: Intermediate × Convergence: Full | 3569 |
| Sampling policy: Exploitative × Convergence: Full | 287 |
| Definition of best target: GP maximum × Convergence: Full | 27 |
| Sampling policy: Intermediate × Definition of best target: GP maximum × Convergence: Full | 2011 |
| Sampling policy: Exploitative × Definition of best target: GP maximum × Convergence: Full | 366 |

The LMM for modelling the difference between algorithm-found and manually defined targets returned an F-statistic of 3.68 and *p*-value of 0.057, with (numerator, denominator) (1, 183) degrees of freedom.
